# CAT tails drive on- and off-ribosome degradation of stalled polypeptides

**DOI:** 10.1101/469296

**Authors:** Cole S. Sitron, Onn Brandman

**Affiliations:** Department of Biochemistry, Stanford University, Stanford CA, 94305

## Abstract

Stalled translation produces incomplete, ribosome-associated polypeptides that Ribosome-associated Quality Control (RQC) targets for degradation via the ubiquitin ligase Ltn1. During this process, the Rqc2 protein and large ribosomal subunit elongate stalled polypeptides with carboxy-terminal alanine and threonine residues (CAT tails). Failure to degrade CAT-tailed proteins disrupts global protein homeostasis, as CAT-tailed proteins aggregate and sequester chaperones. Why cells employ such a potentially toxic process during RQC is unclear. Here, we developed quantitative techniques to assess how CAT tails affect stalled polypeptide degradation in *Saccharomyces cerevisiae*. We found that CAT tails improve Ltn1’s efficiency in targeting structured polypeptides, which are otherwise poor Ltn1 substrates. If Ltn1 fails, CAT tails undergo a backup route of ubiquitylation off the ribosome, mediated by the ubiquitin ligase Hul5. Thus, CAT tails functionalize the carboxy-termini of stalled polypeptides to drive their degradation on and off the ribosome.

---

Stalled mRNA translation produces incomplete polypeptides that can be deleterious to cells. In eukaryotes, Ribosome-associated Quality Control (RQC) recognizes these stalled polypeptides while they are attached to ribosomes and targets them for degradation^1,2^. RQC targets diverse stalled polypeptides generated by a variety of translation abnormalities, including truncated mRNA^3,4^, inefficiently-decoded codons^5^, and translation past stop codons into polyA tails^6,7^.

After recognizing stalls, RQC factors remodel the ribosome to produce a 60S subunit-stalled polypeptide complex^3,4,8–13^ (Fig. 1a). Two proteins, Ltn1 and Rqc2, bind this complex and play central roles in degrading the stalled polypeptide (RQC substrate)^1,2,4,6^. Ltn1, an E3 ubiquitin ligase, ubiquitylates the RQC substrate to mark it for proteasome-mediated degradation^1,2,4,6^. Rqc2 facilitates Ltn1 binding to the ribosome and drives the C-terminal addition of alanine and threonine (“CAT tails”, “CATylation”) to the RQC substrate^2,14–16^. Unlike conventional translation, CATylation occurs without mRNA or the 40S ribosomal subunit^15,16^. Failure to degrade CATylated proteins can result in their aggregation and lead to disruption of global protein homeostasis^17–19^. Understanding why a process as potentially risky as CATylation evolved is a subject of intense interest.

**Figure 1.**
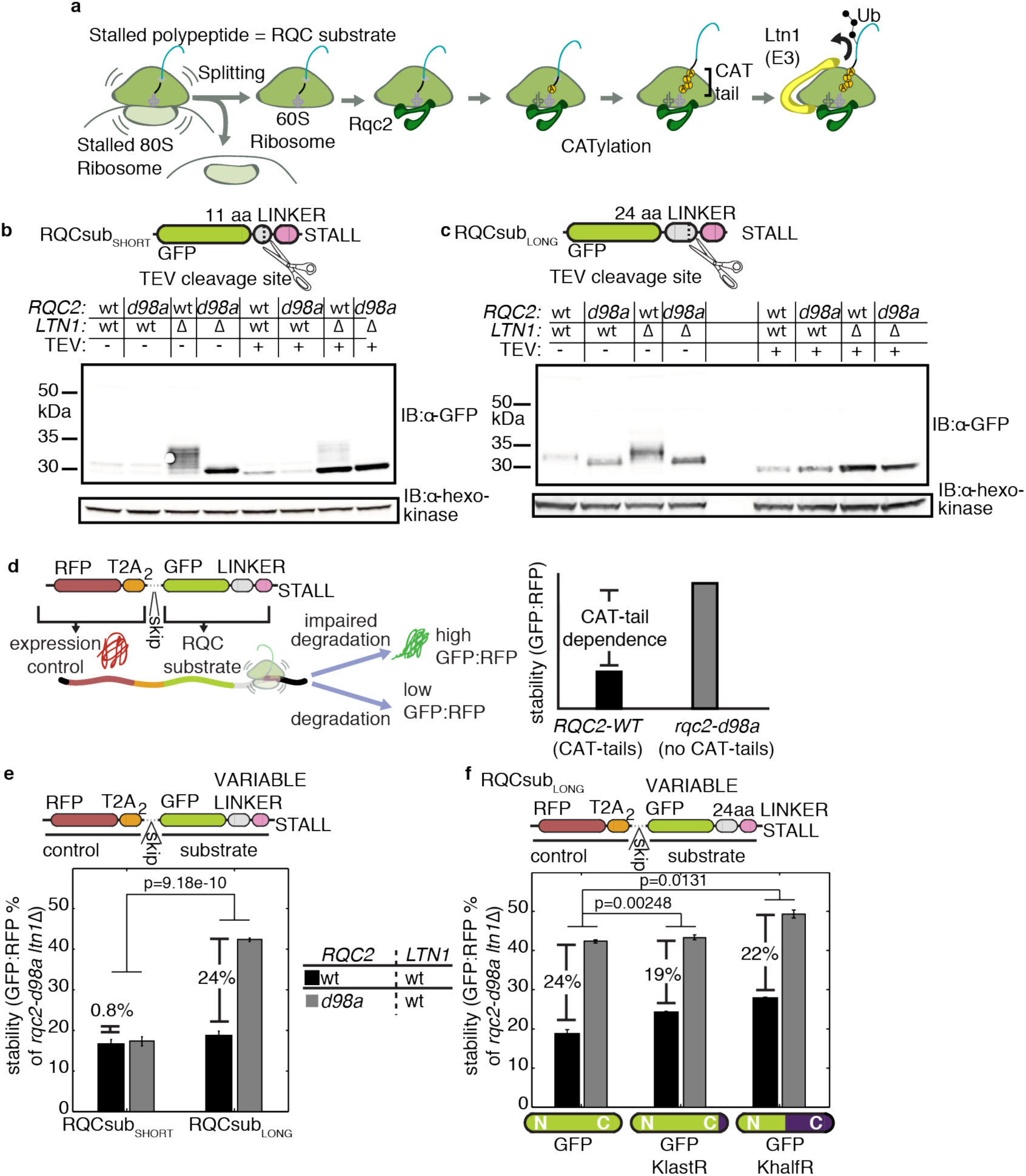
Loss of CAT-tails stabilizes specific RQC substrates. **a**, Model of the RQC pathway. Stalled ribosomes are recognized by a set of factors which facilitate separation of the ribosomal subunits. Rqc2 binds the 60S ribosome-stalled polypeptide complex and directs extension of the stalled polypeptide (RQC substrate) with a CAT tail. Ltn1 binds the complex and ubiquitylates the stalled polypeptide. **b & c**, immunoblots (IBs) of lysates containing two model RQC substrates (schematics above) with and without tobacco etch virus (TEV) protease treatment. GFP, green fluorescent protein. **d**, Schematic of expression-controlled model RQC substrate and definitions of “stability” and “CAT-tail dependence.” RFP, red fluorescent protein. **e & f**, Stability measurements from expression-controlled model RQC substrates with different linker lengths and mutated lysines (0,1, or 10 C-terminal lysines mutated to arginine in GFP). Stability data are reported as means, normalized to the value from the *rqc2-d98a ltn1*Δ strain. CAT tail dependence is indicated with a percent value. Error bars indicate standard error of the mean (s.e.m.) from n = 5 biological replicates and p-values from indicated comparisons of CAT tail dependence are indicated above (16 degrees of freedom, d.o.f.).

In this work, we assessed the contribution of CAT tails to degradation of model RQC substrates using new, quantitative approaches. We found that CATylation enhances ubiquitylation of RQC substrates both on and off the ribosome. On the ribosome, CATylation enhances Ltn1’s ability to ubiquitylate structured substrates, which might otherwise evade Ltn1. Off the ribosome, CAT tails mark escaped RQC substrates for ubiquitylation, dependent on the E3 ubiquitin ligase Hul5.

## CAT tails aid decay of RQC substrates

To measure how CAT tails contribute to RQC substrate degradation, we monitored how disruption of CATylation affects steady-state levels of model RQC substrates in *Saccharomyces cerevisiae*. Each model substrate contained an N-terminal green fluorescent protein (GFP) followed by a flexible linker, tobacco etch virus (TEV) protease cleavage site, and stall-inducing polyarginine sequence (Fig. 1b,c). Ribosomes begin translating these substrates normally but stall within the polyarginine sequence without reaching the stop codon^20^. This produces a GFP-linker-arginine nascent polypeptide that is a substrate for RQC^1^.

We began by constructing two model RQC substrates that differed in the length of the linker (RQCsub_SHORT_, RQCsub_LONG_), allowing two variants for how much of the GFP-linker-arginine polypeptide protruded from the ribosome exit tunnel (Fig. 1b,c). *ltn1*Δ strains accumulated more RQCsub_SHORT_ and RQCsub_LONG_ protein products by SDS-PAGE than *LTN1-WT* strains, confirming that both are substrates for Ltn1 and, thus, RQC (Fig. 1b,c). RQCsub_SHORT_ and RQCsub_LONG_ protein products migrated as higher molecular weight smears in the *ltn1*Δ background when Rqc2 was intact (Fig. 1b,c). These smears collapsed onto discrete bands with mutation of Rqc2 to the CATylation-incompetent *rqc2-d98a* mutant^15^ or cleavage of the C-terminus by TEV protease treatment (Fig. 1b,c), indicating that the smears contained CAT tails of varying length. Strikingly, loss of CATylation with *rqc2-d98a* led to an accumulation of RQCsub_LONG_ but not RQCsub_SHORT_ (Fig. 1b,c). These qualitative results suggest that CAT tails enable efficient degradation of some RQC substrates but not others.

To explore the differences in degradation between RQCsub_SHORT_ and RQCsub_LONG_, we developed a quantitative assay to measure the extent to which RQC substrate degradation depends on CAT tails. We constructed an internal expression control by adding red fluorescent protein (RFP) followed by tandem viral T2A skipping peptides upstream of GFP-linker arginine (Fig. 1d). During each round of translation, the ribosome skips formation of a peptide bond within the T2A sequence, producing an RFP that detaches from GFP-linker-arginine before stalling occurs^21,22^. This detachment ensures that RQC targets GFP-linker-arginine but not RFP. Indeed, *ltn1*Δ did not increase RFP levels compared to wild-type (Extended Data Fig. 1a), confirming that RFP is not an RQC substrate. Because the ribosome synthesizes the two fluorescent proteins stoichiometrically but RQC only targets GFP, the GFP:RFP ratio reports on RQC substrate stability. By comparing stability between an experimental condition and a condition where CATylation and RQC are inactive (*rqc2-d98a ltn1*Δ), we controlled for any RQC-independent degradation or change in fluorescence. This strategy eliminated noise inherent in our model RQC substrate expression system (Extended Data Fig. 1b) and allowed us to quantitatively compare different RQC substrates.

Using our quantitative assay, we re-assessed how degradation of RQCsub_SHORT_ and RQCsub_LONG_ depends on CAT tails. We defined “CAT tail dependence” as the change in protein stability due to CATylation impairment, stability_*rqc2-d98a*_ – stability_*RQC2-wT*_ (Fig. 1d). Consistent with our previous qualitative results (Fig. 1b,c), CAT tail dependence for RQCsub_LONG_ and RQCsub_SHORT_ was 24% and 0.8%, respectively (Fig. 1e). Additionally, we observed substantial CAT tail dependence independent of Ltn1 (in the *ltn1*Δ background) (Extended Data Fig. 1c and subsequent text). To ensure that the CAT tail dependence observed for RQCsub_LONG_ was not an artifact of changes in ribosome stalling, we designed a quantitative stalling reporter similar to one used by the Hegde group^10,23^: *RQC2* or *LTN1* mutations did not affect stalling relative to control (Extended Data Fig. 1d). These data suggest that CAT tails facilitate degradation of RQCsub_LONG_ but are dispensable for RQCsub_SHORT_.

A previous study proposed that CAT tails aid degradation by extending the RQC substrate so that lysine residues buried in the ribosome exit tunnel emerge from the ribosome and can be ubiquitylated by Ltn1^24^. RQCsub_LONG_ has a lysine 24 amino acids away from the stall sequence, placing it in the 35-40 amino-acid-long exit tunnel^25^ at the point of stalling. However, mutation of the buried lysine to arginine (which cannot be ubiquitylated) preserved the bulk of CAT tail dependence (from 24% to 19%) (Fig. 1f; Extended Data Fig. 1e). Similarly, mutation of ten lysine residues in the C-terminal half of RQCsub_LONG_ maintained CAT tail dependence (from 24% to 22%) (Fig. 1f). Although these mutations placed the proximal lysine 150 residues from the stall sequence, *ltn1*Δ still stabilized this substrate (Extended Data Fig. 1e). These data suggest that Ltn1 activity is not restricted to lysine residues close (in primary sequence) to the exit tunnel. Therefore, CAT tails can mediate degradation of RQC substrates without displacing lysine from the exit tunnel.

## Substrate structure affects degradation

We next sought to find the properties of RQCsub_SHORT_ and RQCsub_LONG_ that drive their differences in CAT tail dependence. These substrates differ in their capacity to co-translationally fold (Fig. 2a). After stalling occurs, the linker in RQCsub_LONG_ is long enough to displace ten of the eleven GFP beta strands out of the exit tunnel, which enables the nascent GFP to adopt a stable conformation^26^. By contrast, RQCsub_SHORT_’s linker can only displace nine beta strands, preventing formation of this stable conformation^26^. Thus, a difference in folding states between RQCsub_SHORT_ and RQCsub_LONG_ may account for their different CAT tail dependence.

**Figure 2.**
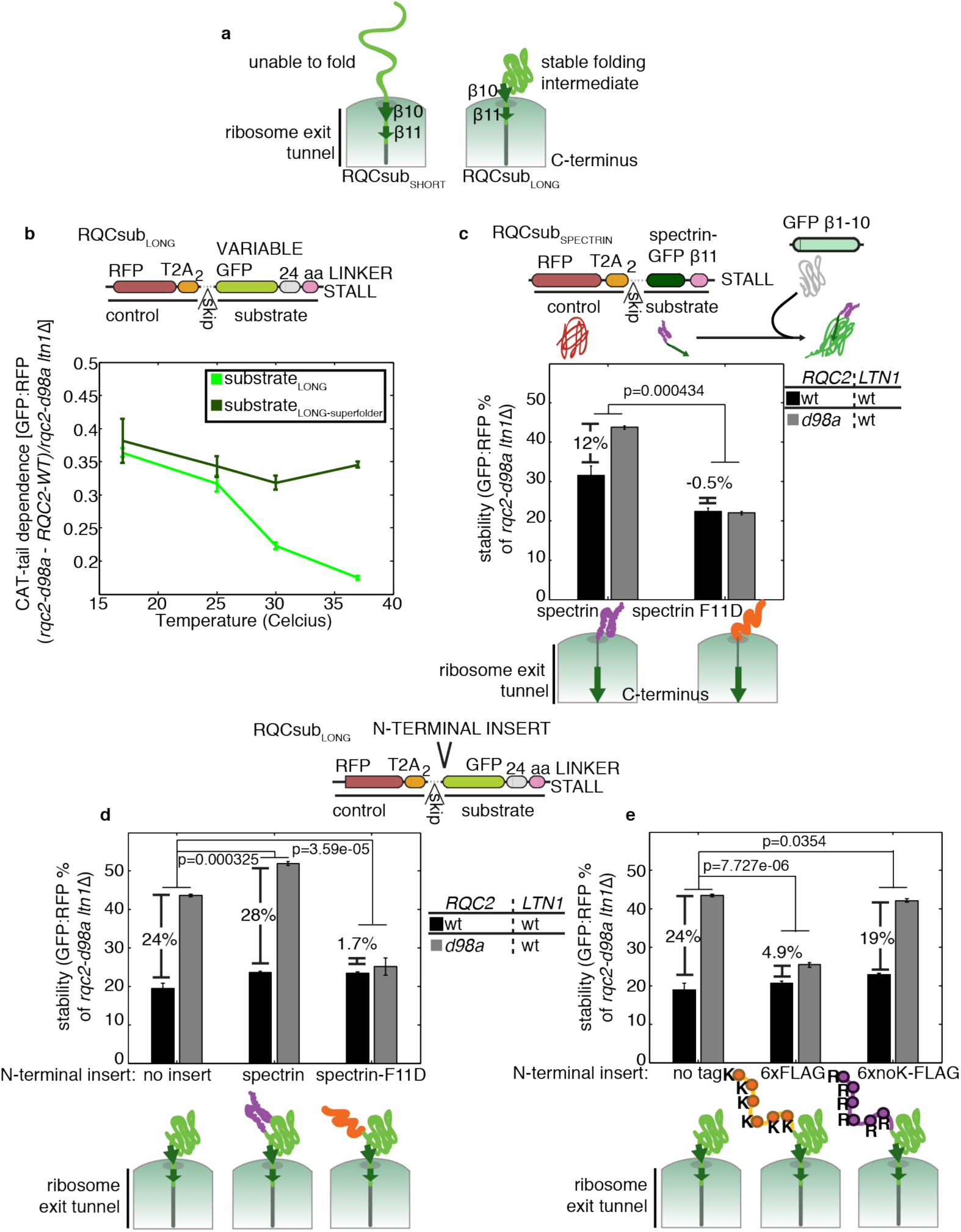
Conditions that favor RQC substrate folding increase CAT-tail dependence. **a**, Cartoon of folding states for RQCsub_SHORT_ and RQCsub_LONG_, emerging from the ribosome exit tunnel. **b**, CAT tail dependence measurements at different incubation temperatures for RQCsub_LONG_ with two different GFP variants (S65T and superfolder). **c**, Normalized stability and CAT tail dependence for a model RQC substrate that can be measured using a split GFP, which features a spectrin R16 domain with or without a fold-disrupting F11D mutation^29^. **d-e**, Normalized stability and CAT tail dependence of variants of RQCsub_LONG_ with added N-terminal spectrin domains (either folded and unfolded spectrin variants) alone or appended to GFP. For all plots, data is normalized as in **Figure 1f** and error bars indicate s.e.m. from n = 3 biological replicates. P values from indicated comparisons are indicated above the bars (8 d.o.f.).

To evaluate the hypothesis that CAT tails promote degradation of structured RQC substrates, we tested how modulating folding of the substrate changes CAT tail dependence. To modulate folding of RQCsub_LONG_, we took advantage of the temperature-sensitive folding of the GFP variant (GFP-S65T) used in the substrate^27^. As incubation temperature increased (decreasing GFP-S65T folding capacity^27^), CAT tail dependence for RQCsub_LONG_ decreased (Fig. 2b). When we replaced GFP-S65T with the less temperature-sensitive superfolder-GFP^27,28^, RQCsub_LONG-superfolder_ had high CAT tail dependence even at high temperatures (Fig. 2b). These data support a model where CAT tails enhance degradation of structured substrates but are dispensable for unstructured substrates.

Next, we tested this hypothesis on spectrin, a protein whose co-translational folding is well-understood and easily modulated^29,30^. We designed RQCsub_SPECTRIN_ using the same RFP-T2A module followed by a spectrin domain, a C-terminal GFP beta strand (β11), and lastly a polyarginine stall sequence. We quantified RQCsub_SPECTRIN_ levels using a “split-GFP” strategy, co-expressing the N-terminal GFP fragment from another transcript^31^. We observed that RQCsub_SPECTRIN_ was 12% CAT tail dependent, while a folding-disrupting mutation^29,30^ eliminated CAT tail dependence (Fig. 2c; Extended Data Fig. 2a). These data suggest that the role of CAT tails in promoting degradation of structured RQC substrates is general, and not unique to GFP.

If CAT tails mediate degradation of structured RQC substrates, we expect that addition of unfolded domains to a structured substrate will relax its CAT tail dependence for degradation. To test this hypothesis, we appended spectrin variants to the N-terminus of RQCsub_LONG_. Folding-disrupted spectrin eliminated CAT tail dependence (from 24% to 1.7%) while folding-competent spectrin did not (from 24% to 28%) (Fig. 2d; Extended Data Fig. 2b). This finding prompted us to inquire whether the presence of unfolded domains alone or, instead, flexible lysine residues within the domains abrogate CAT tail dependence. To distinguish between these possibilities, we added unstructured FLAG-tag variants (with or without lysine residues) to the N-terminus of RQCsub_LONG_. The lysine-containing FLAG-tag decreased CAT tail dependence (from 24% to 4.9%), 3.8-fold more than did lysine-free FLAG-tag (19%) (Fig. 2e; Extended Data Fig. 2c). We thus propose that CAT tails preferentially enhance degradation of RQC substrates lacking lysines in unfolded regions.

## Ltn1-independent ubiquitylation

While impairing CATylation affected the stability of some RQC substrates differently when Ltn1 was intact, impairing CATylation in the *ltn1*Δ background dramatically increased the stability of every substrate we measured (Extended Data. Fig. 1,2). Furthermore, treatment with the proteasome inhibitor bortezomib also increased the stability of RQCsub_LONG_ in *ltn1*Δ, but only when CATylation was intact (Fig. 3a). These data suggest that CAT tails target proteins for proteasomal degradation by Ltn1-dependent and -independent mechanisms.

**Figure 3.**
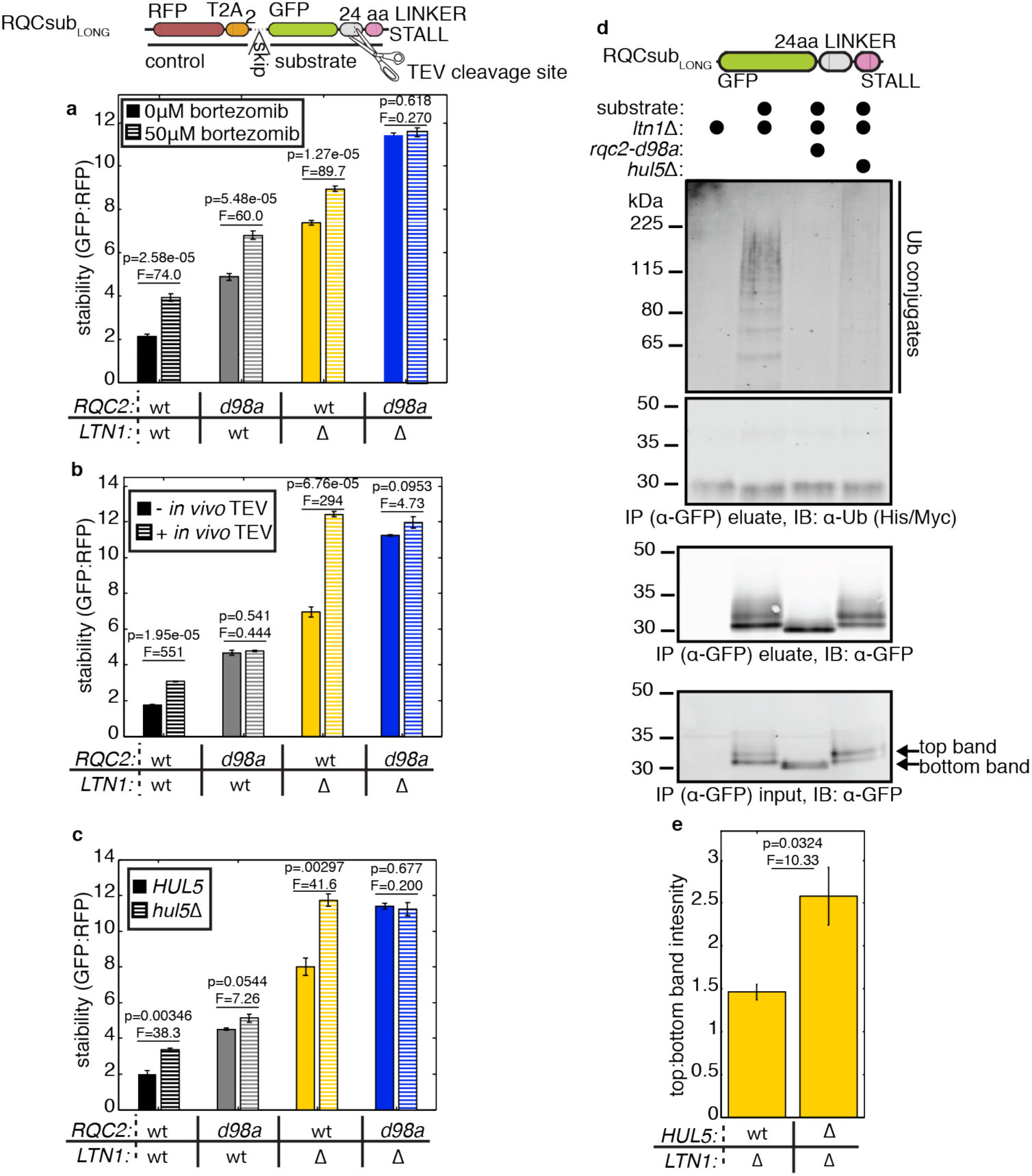
CATylated RQC substrates are ubiquitylated independently of Ltn1. **a-c**, Mean stability of RQCsub_LONG_ in indicated strains after perturbation with the proteasome inhibitor bortezomib, *HUL5* deletion, or TEV protease co-expression (details in panel legend; P- and F-statistics from ANOVA tests with 4 d.o.f. indicated above bars). **d**, Analysis of RQCsub_LONG_ and associated ubiquitin by immunoprecipitation (IP) from indicated cell lysates and IB. **e**, Densitometry analysis of two bands seen in RQCsub_LONG_ immunoblots from whole cell extract, with p- and F-statistics from ANOVA (4 d.o.f.) shown above the bars. Raw images are shown in **Extended Data Fig. 4a**. For all plots, error bars represent s.e.m. from n = 3 biological replicates.

Either the process of CATylation or CAT tails themselves could serve as an Ltn1-independent degradation signal. To distinguish between these possibilities, we employed a strategy to remove a substrate’s CAT tail *in vivo* without disrupting the process of CATylation. We co-expressed RQCsub_LONG_ and TEV protease *in vivo* to cleave RQCsub_LONG_’s C-terminus and remove its CAT tail. TEV co-expression increased RQCsub_LONG_’s mobility on SDS-PAGE (Extended Data Fig. 3a), confirming TEV activity *in vivo*. RQCsub_LONG_ was stabilized by TEV co-expression in cells with Rqc2 intact but not in cells incapable of CATylation (*rqc2-d98a*) (Fig. 3b). Therefore, CAT tails themselves target RQC substrates for Ltn1-independent degradation.

We next searched a set of candidate genes from the ubiquitin-proteasome system to identify an E3 ligase that ubiquitylates CATylated proteins in the absence of Ltn1. Deletion of the proteasome-associated E3 ligase *HUL5^32^* stabilized RQCsub_LONG_ in the *ltn1*Δ background as much as removing CATylation (*rqc2-d98a*) (Extended Data Fig. 3b). Furthermore, during a cycloheximide chase, *hul5*Δ slowed decay of RQCsub_LONG_ in the *ltn1*Δ background as much as impaired CATylation (Extended Data Fig. 3c). This indicates that the cell continuously degrades CATylated proteins when Ltn1 is limiting, dependent on Hul5. The stabilization we observed after loss of Hul5 was not due to perturbed CATylation, as RQCsub_LONG_ had identical amino acid composition in *ltn1Δ and hul5Δ ltn1*Δ (Extended Data Fig. 3d). By microscopy, disruption of *HUL5* did not affect aggregation of RQCsub_LONG_ (data not shown), suggesting that the disruption of degradation did not result from a change in solubility. When Ltn1 was intact, *hul5*Δ specifically stabilized the CATylated species; *hul5*Δ significantly stabilized RQCsub_LONG_ with *RQC2-WT* but not *rqc2-d98a* (Fig. 3c). These modest effects observed in *rqc2-d98a* were likely non-specific, as *hul5*Δ also weakly stabilized a non-stalling degradation sequence (“degron”) (Extended Data Fig. 3e). These results support a role for the E3 ligase Hul5 in Ltn1-independent degradation of CATylated proteins.

Hul5 has E4 ligase activity, which extends existing ubiquitin conjugates to create poly-ubiquitin chains that boost proteasome processivity^32,34,35^. It is thus possible that we identified Hul5 because degradation of CATylated proteins requires extension of a mono-ubiquitin mark left by another E3 ligase. A hallmark of E4 ligase activity is stabilization of the mono-ubiquitylated substrate after loss of the ligase^36,37^, resulting in an 11kDa shift (His-Myc-Ubiquitin) by SDS-PAGE. Purification of RQCsub_LONG_ in the *ltn1*Δ background revealed that *hul5*Δ, like *rqc2-d98a*, diminished detection of ubiquitylated conjugates without stabilizing an apparent mono-ubiquitylated species (Fig. 3d). Thus, Hul5 is required for an E3 ligase activity that ubiquitylates CATylated proteins.

While *hul5*Δ did not stabilize a mono-ubiquitylated species in the *ltn1*Δ background, *hul5*Δ intensified a crisp band within the CATylated smear, ~1kDa above the lowest band (Fig. 3d). This band disappeared after disruption of CATylation (*rqc2-d98a*) (Fig. 3d), suggesting that the corresponding protein contains short CAT tails of relatively uniform size (~10-14 residues). To test whether short CAT tails support Hul5-dependent degradation, we monitored RQCsub_LONG_ stability in the presence of *rqc2-d9a*, an *RQC2* mutant that produces short CAT tails (Extended Data Fig. 4b). In the *ltn1*Δ background, *rqc2-d9a* preserved the majority of RQCsub_LONG_ stabilization after *hul5*Δ that we observed for *RQC2-WT* (Extended Data Fig. 4c). We posit that short CAT tails mark proteins for Hul5-dependent ubiquitylation.

## CAT tails are Hul5-dependent degrons

We wondered whether short tracts of alanine and threonine residues were sufficient to confer the Hul5-dependent degradation we observed for CATylated proteins. To test this, we replaced our model RQC substrates’ stalling sequence with three non-stalling arginine residues (preserving the stalling sequence charge) and appended defined alanine and threonine sequences followed by a stop codon (Fig. 4a). These “hardcoded” CAT tails simulated natural CAT tails but had manipulable sequences and were not RQC substrates. If a hard-coded CAT tail suffices for Hul5-dependent degradation, its stability will be higher in *hul5*Δ cells than wild-type. We define this Hul5-dependence as stability_*hul5*Δ_ – stability_Wt_ (Fig. 4a). While the arginine C-terminus control and alanine/threonine two-mers were not Hul5-dependent, “RRRATA” yielded weak Hul5-dependence (13%) (Fig. 4b). Doubling this motif to form “RRR(ATA)_2_” increased Hul5 dependence to 80%, but the “RRR(ATA)_4_” motif (54%) was weaker than “RRR(ATA)_2_” (Fig. 4b). Therefore, short hard-coded CAT tails suffice for destabilization by Hul5.

**Figure 4.**
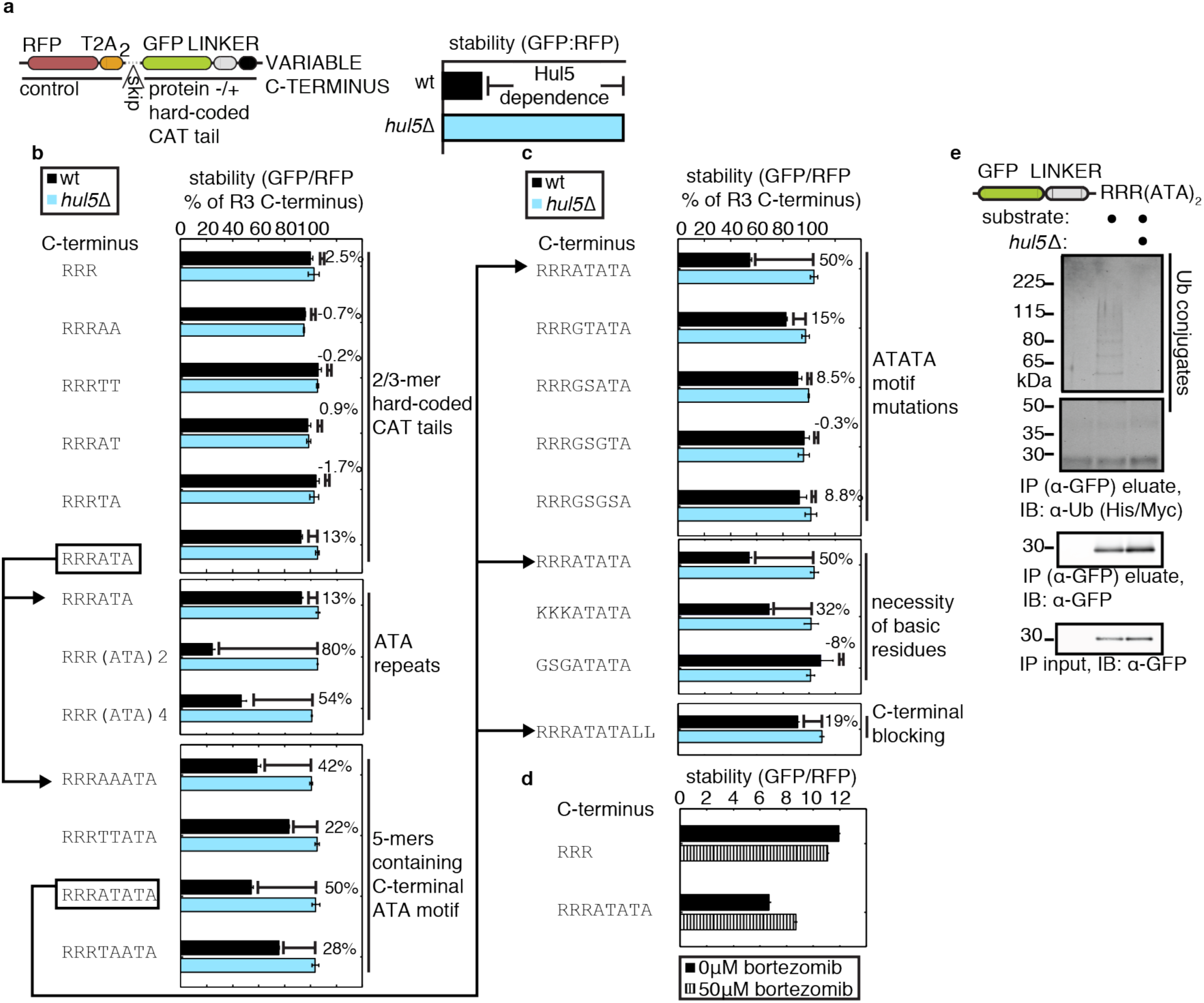
CAT tails are Hul5-dependent degrons. **a**, Schematic of the hard-coded CAT tail construct scaffold, terminating in 3xArg encoded by non-stalling codons (AGA), and definition of Hul5 dependence. **b-c**, Normalized stability measurements of hard-coded CAT tail constructs with indicated C-termini in wild-type and *hul5*Δ cells. Data are presented as mean, normalized to the 3xArg alone construct in wild-type. **d**, Mean stability measurements from wild-type yeast expressing the indicated hard-coded CAT tail constructs, treated with bortezomib. **e**, Analysis of “RRR(ATA)_2_” hard-coded CAT tail construct by IB after IP from lysate. For all plots, error bars represent s.e.m. from n = 3 biological replicates.

We then performed mutagenesis experiments to identify additional CAT tail properties that confer Hul5-dependence. After making modifications to “RRRATATA,” we found that Hul5-dependence decreased after mutating alanine and threonine to glycine and serine (especially alanine adjacent to arginine), replacing arginine residues with non-basic residues, or capping the C-terminus with two leucine residues (a relatively stable C-terminal amino acid^38^) (Fig 4b,4c). This mutagenesis series suggests that CAT tails are effective degrons when: 1) adjacent to basic amino acids, especially when alanine is directly adjacent, and 2) C-terminal.

Given that Hul5 destabilizes hard-coded CAT tails, we investigated whether these proteins are ubiquitylated similarly to naturally CATylated proteins. Stability increased upon bortezomib treatment for “RRRATATA” but not the arginine C-terminus control (Fig. 4d). Purification of the most Hul5-dependent hard-coded CAT tail we tested, “RRR(ATA)_2_” recovered ubiquitin conjugates whose detection was abolished upon *hul5*Δ (Fig. 4e). As for the naturally CATylated RQCsub_LONG_, *hul5*Δ diminished all ubiquitin conjugation and did not stabilize a mono-ubiquitylated species (Fig. 4e). Thus, hard-coded CAT tails are sufficient to mark proteins that are not RQC substrates for Hul5-dependent ubiquitylation. This sufficiency suggests that Hul5-dependent ubiquitylation of RQC substrates can occur off the ribosome, unlike Ltn1-dependent ubiquitylation^39^.

## Dissection of CAT tail function

Our work supports the following model: Ltn1 efficiently ubiquitylates substrates that contain lysine in unstructured regions regardless of whether CATylation takes place (Fig. 5a). However, CATylation enhances Ltn1’s ability to ubiquitylate structured substrates. If Ltn1 fails to ubiquitylate the substrate, short (~1kDa) CAT tails mark that substrate for Hul5-dependent ubiquitylation, which can occur off the ribosome. To support this model, we sought to test its key predictions and quantify how much CAT tails contribute to Hul5 and Ltn1 function.

**Figure 5.**
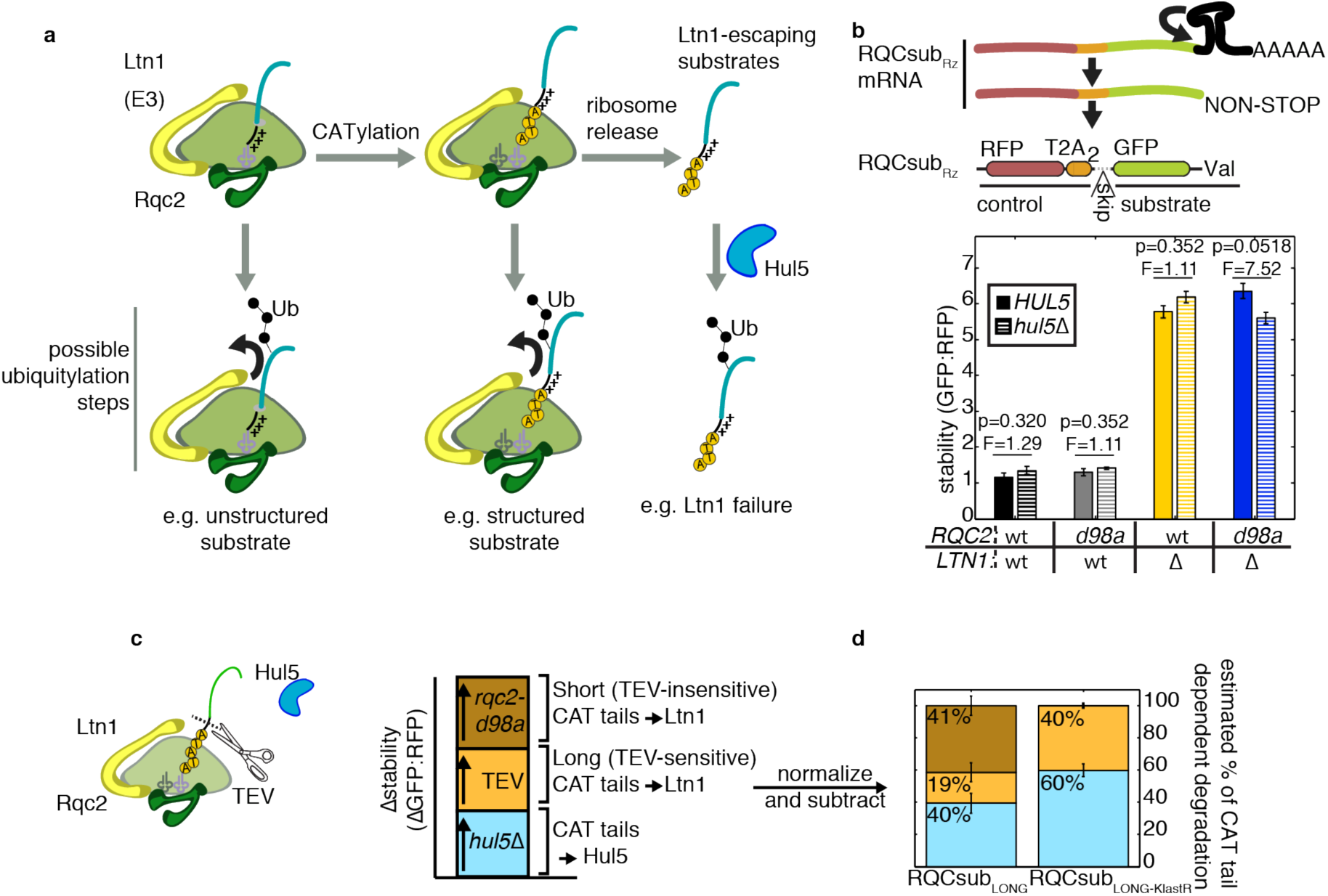
Decomposition of the contribution of CAT tails to Ltn1 and Hul5 function. **a**, Model for how CAT tails enable degradation of RQC substrates by Ltn1 and Hul5. For unstructured substrates, ubiquitylation by Ltn1 occurs efficiently without CAT tails. CAT tails facilitate ubiquitylation of structured substrates, which are otherwise poor Ltn1 substrates. If Ltn1 fails, substrates released from the ribosome can be ubiquitylated, dependent on Hul5. **b**, Mean stability of RQCsub_Rz_ substrate whose mRNA self-cleaves and leaves thus stalls ribosomes without a polybasic tract (see schematic above). Error bars indicate s.e.m. from n = 3 biological replicates and p- and F-statistics from ANOVA (4 d.o.f.) are shown above bars. **c**, Scheme to decompose the contribution of CAT tails to Ltn1 and Hul5 function through combined perturbations to delete *HUL5*, remove long CAT tails with *in vivo* TEV cleavage, then mutate *RQC2*. **d**, Estimation of the the contribution of Ltn1 and Hul5 to CAT tail-mediated degradation of RQCsub_LONG_ and RQCsub_LONG_ with the C-terminal GFP lysine mutated (RQCsub_LONG-KlastR_, as in **Figure 1f**). Data are presented as mean, and error bars indicate s.e.m. from n = 3 biological replicates. Raw data are presented in **Extended Data Figure 5a**.

Our model predicts that CAT tails do not facilitate degradation of substrates that: 1) are unstructured, and 2) terminate in a non-basic residue (preventing Hul5-dependent degradation). We constructed such a substrate, RQCsub_Rz_, encoded by an mRNA containing a hammerhead ribozyme that self-cleaves to produce a non-stop transcript with a truncated 3’ end that stalls ribosomes^40^. Before CATylation, the C-terminal residue of RQCsub_Rz_ is neutrally charged valine and there are too few residues between GFP and the C-terminus to enable formation of the stable GFP conformation^26^. As predicted, neither disruption of CATylation (*rqc2-d98a*) nor loss of Hul5 stabilized RQCsub_Rz_ (Fig. 5b). Thus, unstructured substrates terminating in non-basic amino acids are not CAT tail dependent.

We next revisited RQCsub_LONG_ to dissect how CAT tails mediate its degradation via Hul5 and Ltn1. We first estimated CATylation’s contribution to Hul5 and Ltn1 by measuring the stabilization caused by *hul5*Δ, assuming that Hul5 and Ltn1 activities are independent (Fig. 5c). Using this analysis, we estimated that Hul5 mediates 40% of CAT tail-dependent degradation and Ltn1 mediates the remaining 60% (Fig. 5d). We were additionally interested in analyzing the size of CAT tails that facilitated Ltn1-mediated degradation. To estimate the contribution of long CAT tails, we co-expressed TEV to cleave RQCsub_LONG_ from the ribosome (and evade Ltn1) if its CAT tails were long enough to expose the buried TEV-cleavage-site (greater than 21 residues) (Fig. 5c). TEV co-expression further decomposed Ltn1-mediated degradation into 19% contributed by long CAT tails (TEV-sensitive) and 41% by short CAT tails (TEV-insensitive) (Fig. 5d). We repeated this analysis after mutating RQCsub_LONG_’s exit tunnel-buried lysine. This mutation eliminated the contribution of short CAT tails to Ltn1 function, but increased the relative contributions of CAT tails to Hul5 and long CAT tails to Ltn1 (Fig. 5d). We conclude that CAT tails enable RQC substrates to be targeted by Ltn1 and Hul5. For a structured substrate, short CAT tails may enhance Ltn1 function by exposing lysine residues buried in the exit tunnel. Long CAT tails, some of which may emerge from the exit tunnel, do not require buried lysine in order to support Ltn1 function.

## Conclusions

We propose that CAT tails facilitate ubiquitylation of RQC substrates in two stages. The first stage occurs on the ribosome and is mediated by Ltn1; the second occurs off the ribosome and depends on Hul5.

Our work reveals that the folding states of RQC substrates dictate how cells degrade them. In particular, Ltn1 ubiquitylates structured RQC substrates inefficiently. CATylation enhances Ltn1’s ability to target these substrates, which may arise when translation fails after synthesis of a folding domain (e.g. stalling within polyA when the entire reading frame is synthesized). We propose two models that could explain how CATylation facilitates on-ribosome Ltn1 activity on structured substrates: 1) CATylation impairs substrate folding, or 2) an accessory factor binds CAT tails to promote substrate unfolding or relax Ltn1’s specificity for unstructured substrates. The latter is formally possible because our data suggests that CAT tails long enough to emerge from the ribosome exit tunnel, exposing potential accessory factor binding sites, contribute to Ltn1 function. Future studies will elucidate the mechanisms by which CAT tails enable Ltn1 to ubiquitylate structured substrates.

We discovered that short CAT tails made on the ribosome enable a backup route of RQC substrate ubiquitylation when Ltn1 fails or is limiting. Rather than acting as an inert extension of the RQC substrate, the alanine and threonine residues in CAT tails (along with adjacent basic residues) form a Hul5-dependent degron. Critically, this same alanine and threonine content mediates toxic aggregation of CATylated proteins when the CAT tail is sufficiently long^17–19^. To maximize ubiquitylation by Ltn1 and Hul5 while avoiding aggregation, we speculate that cells coordinate CAT tail synthesis with RQC substrate degradation. How cells accomplish this regulation and ensure CAT tails are of manageable length will be the focus of future research.

## Acknowledgements

We thank S. Marqusee, J. Frydman, Z. Davis, and B. Lu for helpful discussions. We thank L. Steinman, R. Kopito, E.P. Geiduschek, and the members of the Brandman and Kopito Labs for comments on the manuscript. We acknowledge J. Work for his gift of the “10-31” degron and control plasmids. John Schulze performed the Amino Acid Analysis at the UC Davis Genome Center Molecular Structure Facility. Stanford University (O.B.), the US National Institutes of Health (1R01GM115968-01 to O.B.), and the National Institute of General Medical Sciences of the US National Institutes of Health (T32GM007276 to C.S.S.) supported this work.

## Author Contributions

O.B. and C.S.S. conceived of the study and designed the experiments. C.S.S. performed the experiments. O.B. and C.S.S. analyzed and interpreted the data. O.B. and C.S.S. wrote the manuscript. O.B. supervised the study.

## Methods

### Yeast & Growth conditions

Yeast cultures were grown at 30OC (unless otherwise noted) in YPD media or synthetic defined media with appropriate nutrient dropouts.

Deletion strains were constructed in the BY4741 background via transformation with PCR products bearing antibiotic selection cassettes (*NATMX6* or *HYGMX6*). These PCR products contained 40bp of homology to the genome on their 5’ and 3’ ends. Transformants were verified by genomic PCR.

*RQC2* mutants were constructed by first replacing 1.5kb 5’ and 300bp 3’ of the *RQC2* start codon with a *NATMX6* cassette. This gap was repaired in transformants by transformation with a PCR product containing a *LEU2* cassette and *pRQC2-RQC2* variant N-terminus, amplified from plasmids containing these *RQC2* variants.

### TEV treatment

1-liter log-phase yeast cultures were harvested by vacuum filtration and flash frozen in liquid nitrogen before lysis by cryo-grinding. The resulting “grindate” was re-solubilized in TEV buffer (25mM HEPES-KOH pH 7.4, 150mM KOAc, 0.5mM EDTA, 10% glycerol, 0.04% Antifoam-B, and EDTA-Free Pierce Protease Inhibitor Mini Tablets; added 1:1 vol:mass) to produce lysate. The lysate was clarified by centrifugation for 5 min at 5000g, then two rounds of 5 min at 10,000g. 1mM of DTT was freshly added to the lysate before treatment with 2ul ProTEV Plus enzyme (Promega) per 23ul of lysate at 30OC for 3 hr in a thermocycler. 4x NuPage LDS Sample Buffer (Thermo Fisher Scientific) was added at 1:1 vol:vol before boiling at 95OC for 5 min to denature the samples.

### Flow cytometry

Log phase yeast growing in synthetic defined media were measured on a BD Accuri C6 flow cytometer (BD Biosciences) with 3-5 biological replicates (isolated clones) per condition. Data were analyzed using custom Matlab scripts. Plasmid-expressing yeast were selected by gating based on RFP fluorescence. Background signal bleeding from the RFP channel into the GFP channel was calculated using an RFP-only control strain and subtracted before additional calculations. In the case where cultures were treated with bortezomib, treatment lasted for 4 hrs.

### Statistical Analysis

A two-tailed t-test for particular contrast was used to determine whether differences in CAT tail dependence values between substrates were significant. The null hypothesis for the t-test was: μ_*rqc2-d98a* + substrate 1_ – μ_*RQC2-WT* + substrate 1_ = μ_*rqc2-d98a* + substrate 2_ – μ_*RQC2-WT*+ substrate 2_. The linear model was constructed using the raw fluorescence means from each replicate using the “lm” function in R and the linear hypothesis was tested using the “linearHypothesis” function in R.

A one-way ANOVA test (one-tailed) was used to determine whether mean measurements differed between substrates in two different conditions. The null hypothesis was: μ_condition 1_ = μ_condition 2_.

### Immunoprecipitation & Immunoblot

In most cases where results were analyzed by immunoblots or immunoprecipitation, a version of the substrate without the expression-normalizing RFP-T2A module was used. This choice was made to improve clarity by reducing the number of products detected on the gel. The one exception is Extended Data Fig. 3a, where an RFP-T2A-containing substrate was shown in an immunoblot.

For whole-cell immunoblots, 0.375/OD600 ml of yeast culture were pelleted and resuspended in 15ul 4x NuPage LDS Sample Buffer. The sample buffer-resuspended pellets were lysed and denatured by boiling at 95OC for 5 min.

To detect ubiquitin conjugates, cells expressing a plasmid with the bidirectional *pGAL1, 10* promoter driving expression of His-Myc-tagged ubiquitin and the construct of interest were used. An additional plasmid that lacked a construct of interest and only contained His-Myc-tagged ubiquitin was used to assess non-specific ubiquitin detection. A 20ml culture of these cells was grown in SD media overnight containing 1 % galactose and 2% raffinose to induce expression of tagged ubiquitin and the construct of interest. The culture was pelleted by centrifugation, weighed and resuspended in 100mM Tris pH 7.4, 10 mM EDTA at 500ul:25mg pellet. The resuspended culture was added dropwise into liquid nitrogen and cryo-ground. The resultant grindate was resolubilized 1:1 mass:vol in buffer to produce lysate that had a final composition of 50mM Tris pH 7.4, 5mM EDTA, 20mM N-ethylmaleimide (added from a fresh 2M stock in ethanol), 0.5% NP-40, 0.04% NP-40, and EDTA-Free Pierce Protease Inhibitor Mini Tablets. The lysate was clarified by centrifugation at 5000g for 5 min and twice at 10,000g for 5 min. The clarified lysate incubated with equilibrated GFP-Trap magnetic agarose resin (Chromotek) (15ul slurry:25mg pellet) for 1 hr 4OC with rotation, washed 5 times in buffer, and eluted by boiling in 20ul 2x NuPage LDS Sample Buffer per 15ul resin for 95OC for 5 min.

For SDS-PAGE, all samples were run on Novex Nupage 4-12% Bis-Tris gels (Thermo Fisher Scientific) and transferred onto 0.45um nitrocellulose membranes (Bio-Rad) using a Transblot Turbo (Bio-Rad). For ubiquitin detection, the membrane was cut at the 50kDa marker to separate unmodified bands from potential poly-ubiquitylated conjugates; these two halves of the membrane were stained separately. Membranes were blocked for 1 hr with 5% milk in TBST at room temperature before staining with antibodies, either overnight at 4OC or for 4 hr at room temperature. Membranes were stained with the following primary antibodies: 1:2000 Pierce mouse anti-GFP (Thermo Fisher Scientific), 1:1000 rabbit anti-GFP (Life Technologies), 1:3000 rabbit anti-Hexokinase (US Biological), 1:1000 Pierce mouse anti-6xHis (Thermo Fisher Scientific), 1:1000 Pierce mouse anti-Myc (Thermo Fisher Scientific). The following secondary antibodies were used to stain membranes at 1:5000: IRDye 800CW donkey anti-mouse, IRDye 800CW donkey anti-rabbit, IRDye 680RD goat anti-rabbit, or IRDye 680RD goat anti-mouse (LiCor Biosciences). A LiCor Odyssey (LiCor Biosciences) was used to scan immunoblots.

### Amino acid analysis

Yeast lysates were produced as in the “TEV treatment” section, except the following buffer was used to resolubilize the grindate: 50mm HEPES pH7.4, 100mM NaCl, 0.5 mM EDTA. 400ul of clarified lysate was immunoprecipitated 20ul magnetic agarose GFP trap resin. After six washes, the resin was eluted by pipetting up and down in 40ul of buffer adjusted to pH 2.5. Eluates were subjected to amino acid analysis at the UC Davis Genome Center as described in Shen et al^15^.

### Data Availability

The datasets generated during this study are available from the corresponding author upon request.

